# Systems biomedicine of primary and metastatic colorectal cancer reveals potential therapeutic targets

**DOI:** 10.1101/2020.01.25.919415

**Authors:** Mehran Piran, Mehrdad Piran, Neda Sepahi, Ali Ghanbariasad, Amir Rahimi

## Abstract

Colorectal cancer (CRC) is one of the major causes of cancer deaths across the world. Patients survival time at time of diagnosis depends largely on stage of the tumor. Therefore, understanding the molecular mechanisms promoting cancer progression from early stages to high-grade stages is essential for implementing therapeutic approaches. To this end, we performed a unique meta-analysis flowchart by identifying differentially expressed genes (DEGs) between normal, primary and metastatic samples in some test datasets. DEGs were employed to construct a protein-protein interaction (PPI) network. Then, a smaller network containing 39 DEGs were extracted from the PPI network whose nodes expression induction or suppression alone or in combination with each other would inhibit tumor progression or metastasis. A number of these DEGs were then verified by gene expression profiling, survival analysis and a number of validation datasets from different genomic repositories. They were involved in cell proliferation, energy production under hypoxic conditions, epithelial to mesenchymal transition (EMT) and angiogenesis. Multiple combination targeting of these DEGs were proposed to have high potential in preventing cancer progression. Some genes were also presented as diagnostic biomarkers for colorectal cancer. Finally, TMEM131, DARS and SORD genes were identified in this study which had never been associated with any kind of cancer neither as a biomarker nor curative target.

## 1. Introduction

Colorectal cancer (CRC) is a major global medical burden worldwide (1). Approximately more than one million people are diagnosed with CRC each year, and about half of them die of CRC annually (2). Complex genetic interactions are combined with environmental factors to trigger a cell to become cancerous. Among them, aberrant growth factor signals contribute to uncontrolled proliferation of cells which ultimately lead to metastasis. Contrary to early-stage tumor cells, malignant cells have the ability to detach from the stroma as well as acquire motility (3). This event happens during a process called EMT in which cells lose their epithelial characteristic including adhesion and subsequently dedifferentiate into mesenchymal mobile cells (4). Therefore, Investigating DEGs between primary and metastatic sites of tumors would aim to recognize key factors playing role in cell migration. We performed the statistical analysis between primary sites and metastatic sites in one part of the analyses. While primary sites were either benign or malignant colon biopsies, metastatic sites were located on the other organs.

A large number of molecular and pathway targets have been identified for treatment of CRC during the past decades. Besides, growing progresses have been made in development of chemotherapy and antibody drugs (5). Tyrosine kinase (TK) targeting using monoclonal antibodies and small molecule tyrosine kinase inhibitors are effective strategies (6). Targeting cancer-related inflammation biomarkers like IL-6/JAK/STAT3 pathway which inhibits progression of solid tumors is another beneficial therapeutic strategy (7). In addition, restraint of cytosolic β-catenin via disturbing hyperactive Wnt/β-catenin signaling pathway could be another treatment approach for colorectal and many other types of cancer (8). Inhibition of matrix metalloproteinases (MMPs) and TGFβ signaling pathways is a therapeutic approach to prevent liver metastasis (9) (10, 11). Furthermore, PI3K inhibition suppresses lung metastasis in CRC patients (12, 13). Among the known anticancer drugs, Cetuximab is one of the popular ones which is a monoclonal antibody against epidermal growth factor receptor (EGFR) (14). Furthermore, vascular endothelial growth factor (VEGF) antibody, bevacizumab, is the standard treatment for metastatic colorectal cancer (15).

One way to identify molecular mechanism of pathogenesis in a biological context is to analyze transcriptomic data. In this study, we conducted a unique meta-analysis flowchart which contained three datasets for retrieving differentially expressed genes considered as the test set and seven datasets to validate selected DEGs considered as the validation set. The test datasets were constructed from three microarray studies and DEGs were excavated for any pairwise comparison between four groups of samples. Common DEGs between similar analyses in test datasets were regarded as final DEGs employed for PPI network assembly. With more datasets, few common DEGs were found with the desired log-fold change and p-value thresholds which were not sufficient for network construction. 12 common genes called core genes were recognized that their expression is different between primary and metastatic sites. Then, a smaller network called core network was extracted from the PPI based on a shortest-path based scoring algorithm containing core genes and 27 neighboring genes. To compensate for the small number of datasets in test set, validation datasets were employed from different genomic repositories to validate selected DEGs in the core network (see Figure 1). Besides, expression profiling and survival analysis provided more evidence about the accuracy of our results. We obtained some DEGs involved in cancer progression whose expression could be targeted (suppressed or induced) individually or in combination with one another for CRC treatment. Moreover, Some of those gene expressions were proposed to be CRC biomarkers.

**Figure 1.**
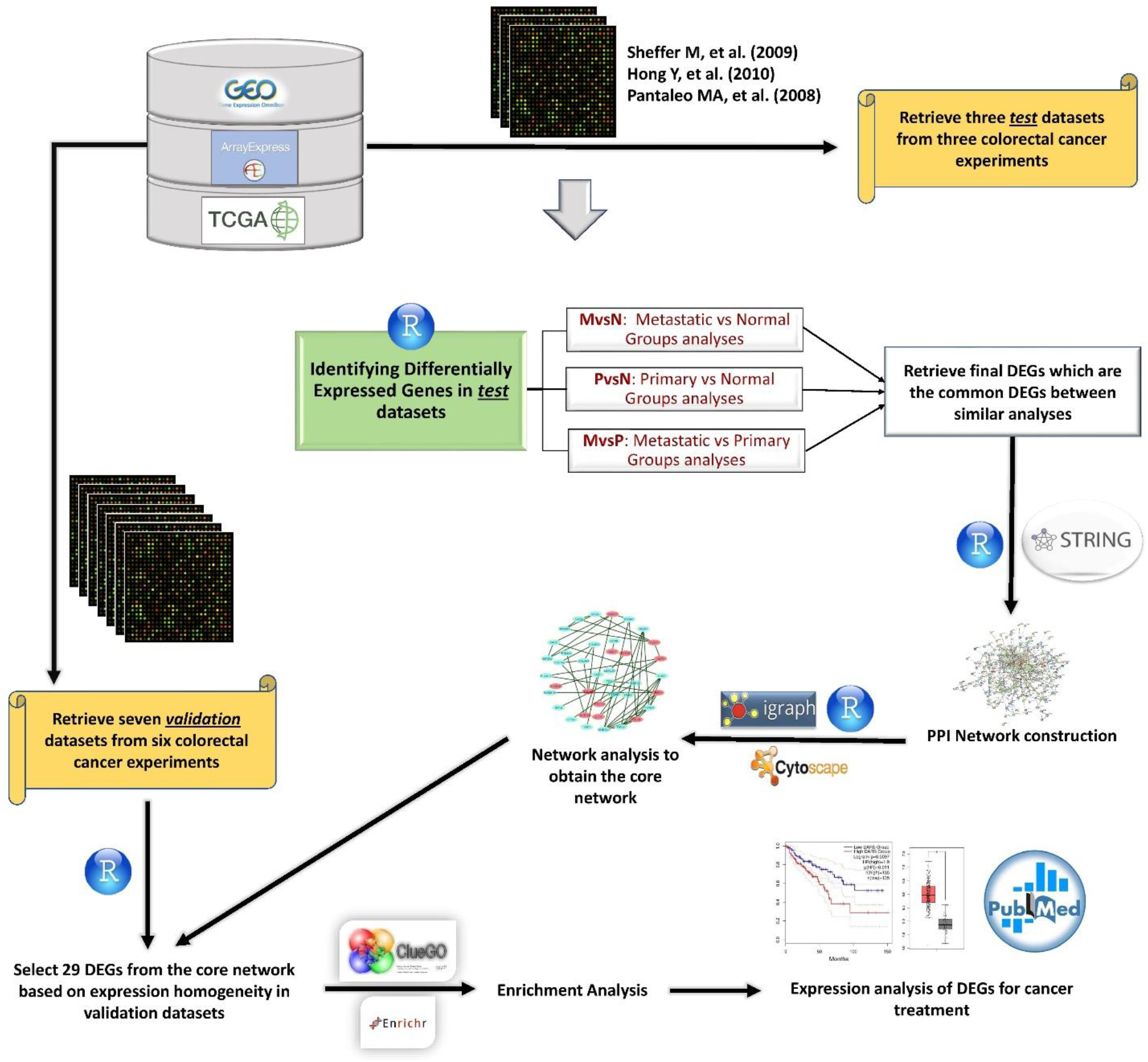
The meta-analysis flowchart to attain therapeutic genetic targets. Gene expression datasets were extracted from different databases. Data were analyzed and visualized using R programming language. DEGs were obtained from analyzing test datasets which then were verified by the validation datasets. STRING database was utilized to construct the PPI network from DEGs, R software was used to analyze the network, Cytoscape was employed to visualize the networks and enrichment results were obtained from ClueGO Cytoscape plugin and Enrichr online tools. Next, survival analysis and expression profiling were used for more validation of expression results. Finally, our results were compared to other studies and molecular mechanism of validated DEGs were interrogated to propose a combination of target therapies.

## 2. Materials and Methods

### 2.1 Database Searching and recognizing pertinent experiments

Gene Expression Omnibus (GEO) (http://www.ncbi.nlm.nih.gov/geo/) and ArrayExpress (https://www.ebi.ac.uk/arrayexpress/) repositories were searched to detect test set experiments containing high-quality transcriptomic samples concordance to the study design. Colorectal/colon cancer, primary, EMT and metastasis are the keywords utilized in the search and it was filtered for Homo sapiens. Microarray raw data with accession numbers GSE41258, GSE9348 and GSE10961 were downloaded from GEO and ArrayExpress. Two datasets were assembled from samples in GSE41258 study. One contained liver metastasis, primary and normal samples and another contained lung metastasis, primary and normal samples. Third dataset was constructed from two microarray studies. Normal samples and primary samples were extracted from GSE9348 study and liver metastatic samples were obtained from GSE10961 study. In all datasets, Normal samples were healthy colon tissues adjacent to the primary tumors and primary samples were in stage I and II (non-metastatic) of colorectal cancer. To make the validation datasets, two new datasets were constructed from GSE41258 study whose samples were not present in the test datasets one contained liver metastasis and another contained lung metastasis samples. A dataset was constructed from count RNAseq files in The Cancer Genome Atlas (TCGA) database (TCGA dataset). Three RNAseq datasets were constructed from GSE50760, GSE144259 and GSE89393 studies encompassing liver metastatic samples. The last dataset was built from GSE40367 microarray study containing liver metastasis samples. Some primary groups in validation datasets were metastatic.

### 2.2 Identifying Differential Expressed Genes in Microarray Datasets

R programming language (v3.6.2) was used to import and analyze data for each dataset separately. Preprocessing step involving background correction and probe summarization was done using RMA method in “affy” package (16). Absent probesets were identified using “mas5calls” function in this package. If a probeset contained more than two absent values in each group of samples, that probeset was regarded as absent and removed from the expression matrix. Besides, Outlier samples were identified and removed using PCA and hierarchical clustering methods. Next, data were normalized using Quantile normalization approach (17). Then, standard deviation (SD) for each gene was computed and median of all SDs was utilized as a cutoff to remove low-variant genes. Therefore, low-variant genes no longer influenced the significance of the high-variant genes. Many to Many problem (18) which is mapping multiple probesets to the same gene symbol was solved using nsFilter function in “genefilter” package (19). This function selects the probeset with the highest Interquartile range (IQR) to map to the gene symbol. “LIMMA” R package which applies linear models on the expression matrix was utilized to discover DEGs between all groups of samples (20). Genes with absolute log fold change larger than 0.5 and Benjamini-Hochberg adjusted p-value (21) less than 0.05 were selected as the DEGs.

### 2.3 Identifying Differential Expressed Genes in RNAseq Datasets

Count files for five primary samples containing more than 90 percent tumor cells as well as five normal samples involving 100 percent normal cells were downloaded from TCGA database. They were imported into R and merged together to construct the TCGA dataset. Then, using “DESeq2” R package (22) data were normalized and DEGs were identified between two groups. For RNAseq datasets in GEO, FPKM normalized data were downloaded and imported into R. data were log-2 transformed and using “LIMMA” R package, DEGs were identified between the groups.

### 2.4 Network Construction

STRING database was used to generate Protein-Protein Interaction (PPI) network. Final DEGs were given to database and sources of evidence were chosen to create interactions. Interactions were downloaded and imported into R. Next, nodes and interactions were obtained separately using “igraph” package in R software (23) and giant component of the weighted network was extracted from the whole network. Next, the weighted adjacency matrix of the giant component was transformed into a symmetric matrix to get modified into a new adjacency matrix using topological overlapping measure (TOM) function in “WGCNA” R package (24). To remain with a distance matrix, the new adjacency matrix was subtracted from one.

### 2.5 Neighbourhood Ranking to the Core Genes

Using R software, a matrix of all shortest paths, called SP, between all pairs of nodes for the weighted network was constructed using Dijkstra algorithm (25). By utilizing this matrix, a distance score, Dj, for each node in the PPI network was computed. Dj is the average of the shortest paths from all the non-core genes to reach the node j subtracted from the average of the shortest paths from the core genes to reach the node j divided by the average of the all shortest paths to reach the node j from the whole network. This scoring system implies how much close each node is to the core nodes (26, 27).

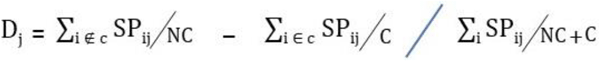

C is the number of core nodes and NC is the number of non-core nodes. ∑_i_ ∉ c SP_ij_ is the sum of all distances, in SP, between node j and all the non-core nodes. ∑_i_ ∈ c SP_ij_ is the sum of the distances between node j and all the core nodes. ∑_i_ SP_ij_ is the sum of the distances between node j and all the nodes. A positive score implies that node j is closer to the core nodes compare to the rest of the nodes. Nodes with positive scores were kept and the others were removed from the network. It should be noted that D scores were calculated without imposing any threshold on edge weights.

### 2.6 Enrichment Analysis

Enrichment analysis was performed using ClueGO Cytoscape plugin (28). Enriched terms for biological processes were obtained from GO repository. For pathway enrichment analysis, information in KEGG (29), Reactome (30) and WikiPathways (31) databases were used. P-value were adjusted using Benjamini-Hochberg method and cut off was set on 0.05. In addition to Cytoscape, Enrichment analysis was performed using Enrichr online tool (32) as well. Enriched terms for biological processes were obtained from GO repository. For pathway enrichment analysis, WikiPathways signaling repository version 2019 for humans was used. Enriched terms with the top score and the p-value less than 0.05 were selected.

### 2.7 Analyzing Gene Expression Profiles

Genes were given to GEPIA2 web server to validate identified DEGs based on datasets in TCGA genomic database (33, 34). For boxplots, expression profiles were compared between tumor and normal samples in multiple colorectal adenocarcinoma (COAD) datasets. LogFC cutoff was 0.5 and p-value was 0.01. TPM normalized data were log2 transformed. For survival plots, Overall Survival option was selected and median was chosen to define the border of High and Low groups. 95% confidence interval was set for analysis. All COAD datasets with monthly expression values were selected in order to obtain survival results.

## 3. Results

### 3.1 Data Preprocessing in Test Datasets

Each dataset was imported into R separately. outlier sample detection was conducted using PCA and hierarchical clustering approaches. Figure2A illustrates the PCA plot for samples in the first dataset. The same plot was created for the second and third datasets. Between the three groups of samples in each dataset, a number of them were separated from their sets along the PC axes and they were considered as outliers. To be more specific, a hierarchical clustering method introduced by Oldham MC, et al (35) was used. To compute the distances between samples, Pearson correlation analysis was performed between them and coefficients were subtracted from one. Figure 2B depicts the dendrogram for normal samples. In Figure 2C normal samples were plotted based on their Number-SD scores. To get this number for each sample, the average of whole distances was subtracted from the distance average of each sample. Then, results of these subtractions were normalized by standard deviation of sample distance averages (35). Samples with Number-SD less than negative two are usually far apart from their cluster set in the PCA plane and they were regarded as outliers in our analyses. sixteen outlier samples for GSE41258 test datasets and three outliers for the third dataset were recognized. Supplementary file1 contains information about the groups of samples.

**Figure 2.**
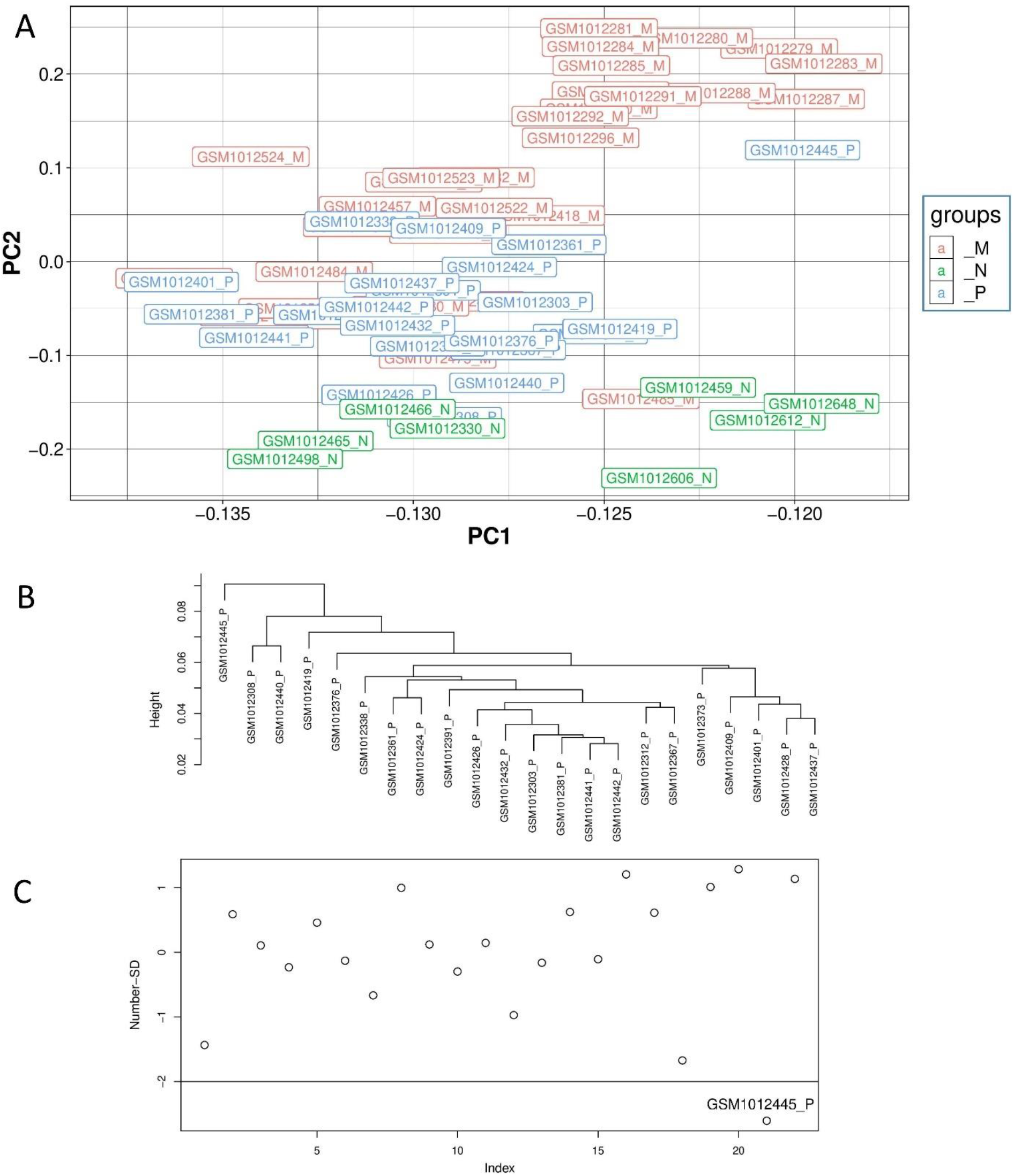
Illustration of outlier samples in the first dataset. A, is the PCA plot, B is the dendrogram for the primary samples and C is the Number-SD plot for primary samples. GSM1012445 is an outlier sample in primary group as it has located in a distance from its group in the PCA plane. In addition, it has formed a unique cluster in the dendrogram and its Number-SD score is less than negative two.

Supplementary file 2 illustrates the average expression values for some housekeeping genes and DEGs between Primary and Metastatic samples. Common DEGs between lung-primary analysis and liver-primary analysis with absolute LogFC larger than 1 in GSE41258 datasets were illustrated in A. The same plot was made for the common DEGs in liver-primary analysis in the third dataset in B. Housekeeping genes were situated on the diagonal of the plots whilst DEGs were located above or under the diagonal. This demonstrates that the preprocessed data was of sufficient quality for downstream analysis.

### 3.2 Meta-Analysis and Identifying Differentially Expressed Genes

10891 unique DEGs with adjusted p-value < 0.05 and absolute log fold change > 5 were achieved from eight groups of DEGs yielded from eight independent analyses on three test datasets. They included two analyses of liver metastasis versus normal, two analyses of liver metastasis versus primary, one analysis of lung metastasis versus normal, one lung metastasis versus primary and two analyses of primary versus normal (Table 1). Liver metastasis contained metastatic colorectal samples taken out from liver tissue and lung metastasis involved metastatic colorectal samples obtained from lung tissue. Note that the majority of primary groups in validation datasets were metastatic. In fact, This kind of sample selection provided some DEGs for us that could significantly contribute to progression of tumor not only in the primary site but also to liver and lung tissues. Common DEGs between all metastasis vs normal analyses were 155 genes. Common DEGs between all metastasis vs primary analyses were 72 genes. Common DEGs between all primary vs normal analyses were 239 genes. There were 334 unique DEGs between these three sets of analyses. Finally, from these 334 DEGs, 242 of them were identified to be in all test and validation analyses considered as the final DEGs. There were 12 final DEGs in primary versus normal analyses considered as the core genes. All DEG sets and their LogFC are presented in Supplementary file 3.

**Table 1.** The practical information for the core network DEGs in test dataset. Up means gene was upregulated and Down means gene was Downregulated. MvsN contains all the metastatic versus normal analyses, PvsN contains all the primary versus normal analyses and MvsP contains all the metastatic versus primary analyses. Some genes are present in more than one analysis.

| MvsN |  |  |  | BvsN |  |  |
| --- | --- | --- | --- | --- | --- | --- |
| DEGs | GSE41258 |  | GSE9348_GSE10961 | DEGs | GSE41258 | GSE9348_GSE10961 |
|  | liver-normal | lung-normal | liver-normal |  | primary-normal |  |
| ETHE1 | Down | Down | Down | SGK1 | Down | Down |
| DARS | Up | Up | Up | EIF2S2 | Up | Up |
| TMEM131 | Down | Down | Down | TRIB3 | Up | Up |
| TST | Down | Down | Down | DARS | Up | Up |
| LGALS4 | Down | Down | Down | RFC3 | Up | Up |
| TRIB3 | Up | Up | Up | ETHE1 | Down | Down |
| COL5A2 | Up | Up | Up | TOP2A | Up | Up |
| COL4A1 | Up | Up | Up | CKS2 | Up | Up |
| PTP4A1 | Down | Down | Down | SORD | Up | Up |
| COL1A2 | Up | Up | Up | PSMA7 | Up | Up |
| SORD | Up | Up | Up | GPX3 | Down | Down |
| SQOR | Down | Down | Down | MAD2L1 | Up | Up |
| OAS1 | Down | Down | Down | SQOR | Down | Down |
| TRIM31 | Down | Down | Down | TST | Down | Down |
| TWF1 | Down | Down | Down | GINS1 | Up | Up |
| MvsP |  |  |  | LGALS4 | Down | Down |
| DEGs | GSE41258 |  | GSE9348_GSE10961 | PTP4A1 | Down | Down |
|  | liver-primary | lung-primary | liver-primary | PLAGL2 | Up | Up |
| ETHE1 | Down | Down | Down | COL1A2 | Up | Up |
| ATF5 | Up | Up | Up | CTSH | Up | Up |
| CDC6 | Down | Down | Down | MT2A | Down | Down |
| TMPO | Down | Down | Down | COL5A2 | Up | Up |
| P4HA1 | Up | Up | Up |  |  |  |

### 3.3 Undirected Protein-Protein Interaction Network

242 final DEGs were utilized to construct the Protein-Protein-Interaction (PPI) network. STRING database was employed to generate the Interactions based on seven sources of evidence namely, Neighborhood, Text mining, Experiments, Databases, Co-expression, Gene fusion and Co-occurrence. STRING combined scores were used as the edge weights. The giant component of the weighted network with 205 nodes and 554 edges was depicted in Figure 3.

**Figure 3.**
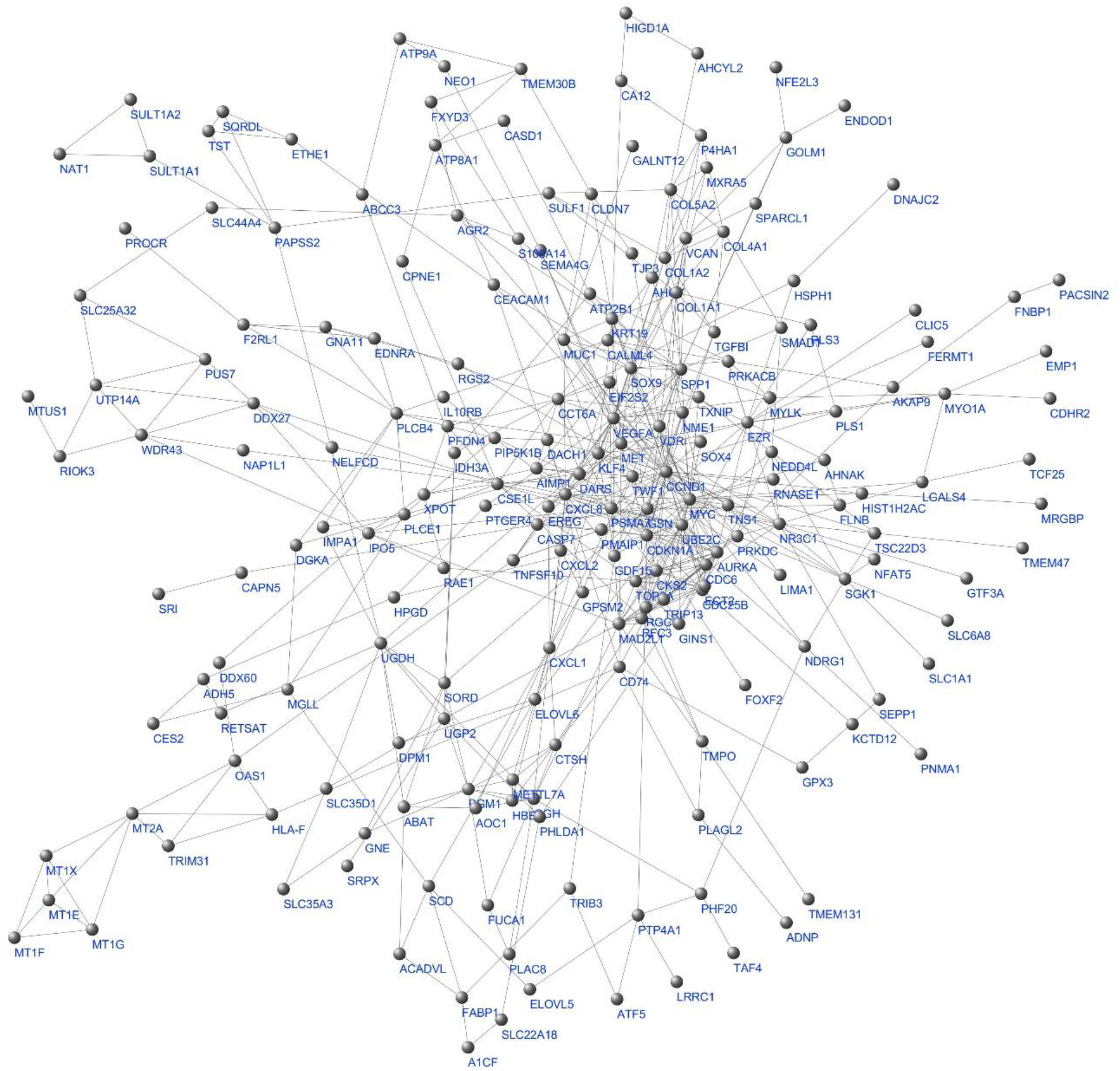
The whole network giant component. Labels are protein/gene symbols. This is a scale free network (116) which follows a power law distribution (most of the network nodes have a low degree while there are few nodes with high degree).

### 3.4 Determination of Core Genes Neighborhood through Shortest Path-Based Scoring System

In this step, interactions combine scores which came from all sources of evidence in STRING database were converted to weights between nodes and these weights were used as the estimation of distances in the weighted adjacency matrix. Nodes with shorter distances from the core genes were selected and a smaller network was extracted from the main network. Computing the shortest path score for the non-core genes led to a network of 39 nodes comprising 12 core nodes and 27 neighbors. This multi-component graph called core network illustrated in Figure 4.

**Figure 4.**
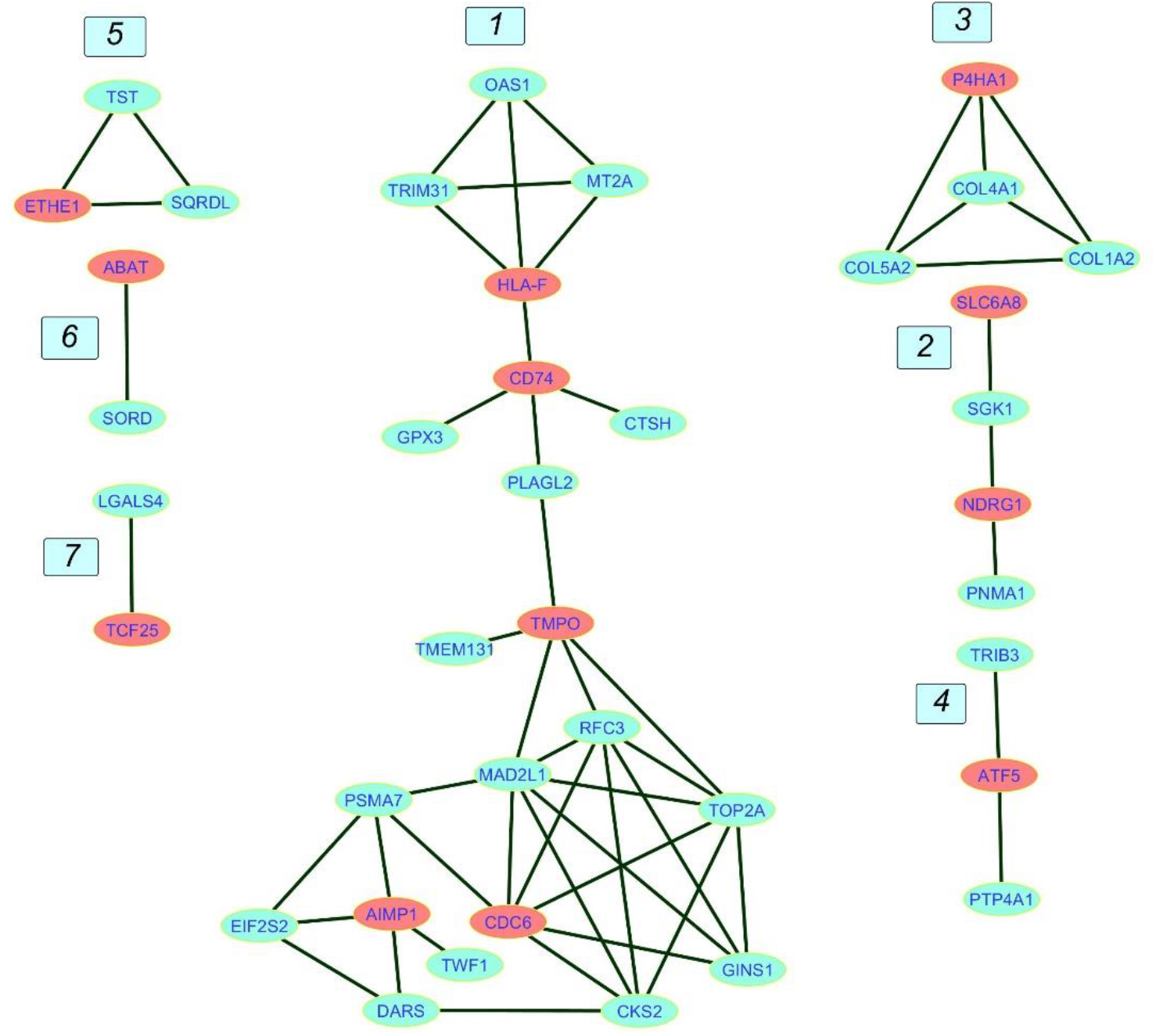
The Core network. The network contains seven components numbered from 1 to 7. Component 1 is the giant component. core genes are in red and non-core genes are in blue.

Majority of the nodes in core network were selected for investigation based on the similarity of expression patterns in all datasets. Expression states of selected genes between any pair-wise comparisons were depicted in Table1. For the three Metastatic-Normal comparisons (MvsN), most of the nodes exhibited similar expression pattern. The same was true for all Primary-Normal analyses (PvsN) and Metastatic-Primary comparisons (MvsP). Heatmaps were illustrated in Figure 5 for all members of the core network in three datasets. Clustering was performed by applying Euclidean distance and complete method on gene expression values. Genes present in the top right corner of the three plots possessed high expression values in colon tissues. Moving from border to the center of plots, we go from Normal to primary and from primary to metastatic samples. Some genes exhibited a descending expression trend such as mitochondrial genes ETHE1, TST and SQOR. Few genes witnessed an ascending trend such as collagen genes and SORD and P4HA1.

**Figure 5:**
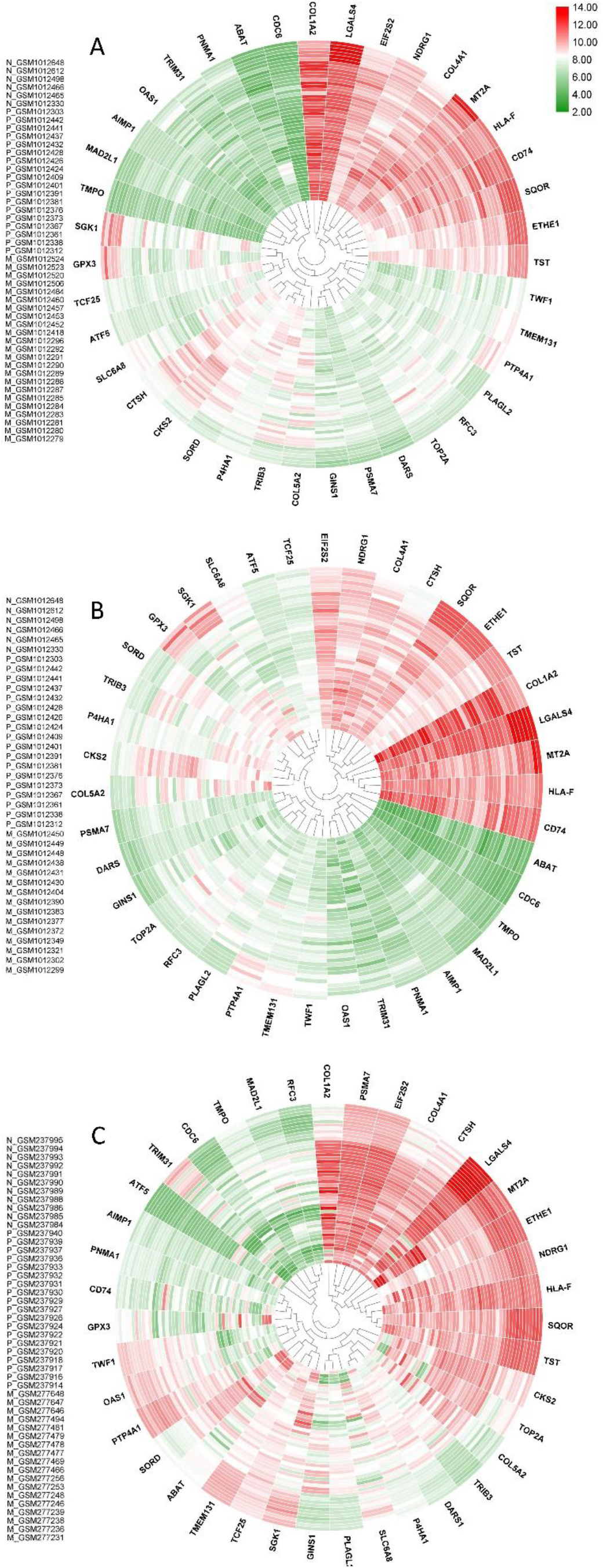
Core network genes heatmaps. A is dataset1, B in dataset2 and C is dataset3. Values were obtained from expression matrices, Log2 transformed and normalized by quantile method. Sample IDs are placed in the left hand side of each plot. Normal samples start with N, Primary samples start with P and liver and lung metastatic samples start with M. Outer samples are normal, middle samples are primary and inner samples are metastatic. Genes were clustered together based on hierarchical clustering.

### 3.5 Network Descriptive

The giant component diameter was eight containing TRIM31, HLA-F, CD74, PLAGL2, TMPO, MAD2L1 PSMA7, AIMP1 and TWF1. Transitivity was around 60%, edge density was about 18% and the mean distance was 3.48. Two important centralities, Degree and Betweenness, along with other centralities and the average distances for giant component nodes are provided in Supplementary file 4. MAD2L1 had the highest Degree and a relatively high betweenness. TMPO had the highest Betweenness and a pretty high degree. Similar to TMPO, its direct neighbor, PLAGL2 had a rather high Betweenness. This gene linked the two parts of the giant component together.

### 3.6 Processing Validation Datasets

Core network DEGs were identified in the seven validation datasets. They were presented in Table 2. Most of the DEGs illustrated similar results in both Table1 and Table2 which proves the accuracy of obtained DEGs from test datasets. Expressions of genes that were totally homogeneous in each of MvsN or MvsP or PvsN analyses are presented in green and the ones that differed only in one analysis are presented in yellow. Expression of Genes that were different in more than one dataset are in white. DEGs with Absolute LogFCs less than 0.2 were not reported in Table2. Expression analysis of all validation datasets are presented in Supplementary file 3.

**Table 2.**
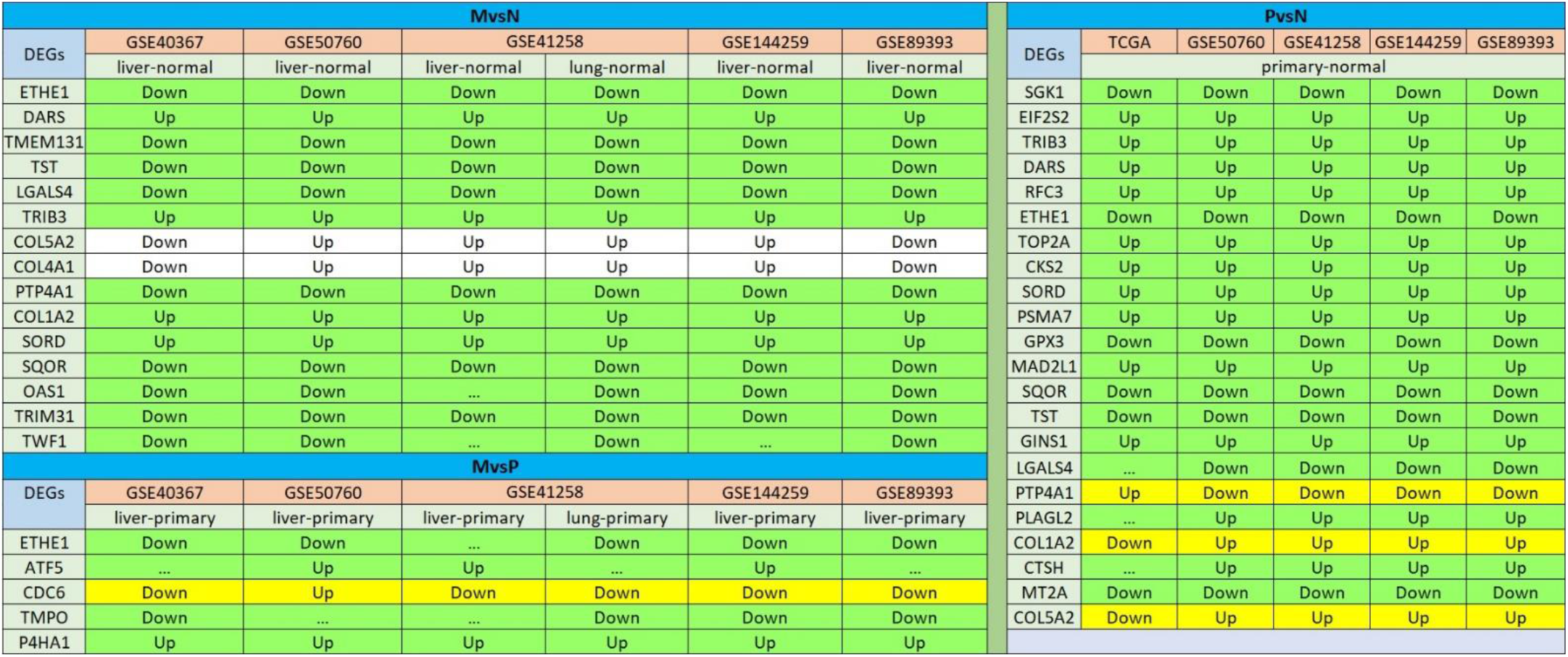
Illustration of core network DEGs in validation datasets. Up means gene was upregulated and Down means gene was Downregulated. MvsN contains all the metastatic versus normal analyses, PvsN contains all the primary versus normal analyses and MvsP contains all the metastatic versus primary analyses. Some genes are present in more than one analysis. The expression status for the genes in green rows are similar in all datasets regardless of empty cells. In the yellow rows only one dataset is different from the others and in the white rows genes exhibited a heterogeneous expression status in different datasets.

### 3.7 Enrichment Analysis

Figure 6 illustrates the enrichment results for the core network genes using ClueGO software. Three signaling databases called KEGG, Reactome and WikiPathways were used for the pathway enrichment. Biological Function terms were enriched from GO database. Genes and terms associated to a specific cellular mechanism formed distinct components. Different pathway terms related to polymerization and degradation of collagens in extra cellular matrix (ECM) has emerged in blue component of enrichment graph which formed a distinct component (component 3) in the core network. In tumor environment different concentrations of collagen fibers are regularly secreted and degraded and aligned together to create ECM stiffness suitable for cellular migration (36, 37). Genes that were enriched for sulfide oxidation terms formed a distinct component in the core network as well. Genes in green are engaged in interferon gamma signaling that has dual roles in cancer. On the one hand, INF-γ has anti-proliferative roles by employing different mechanisms such as induction of p21 (38), induction of autophagy (39), regulation of EGFR/Erk1/2 and Wnt/β-catenin signaling (40) and so on. On the other hand, it enhances the outgrowth of tumor cells with immunoevasive properties depending on cellular and microenvironmental context (41, 42).

**Figure 6.**
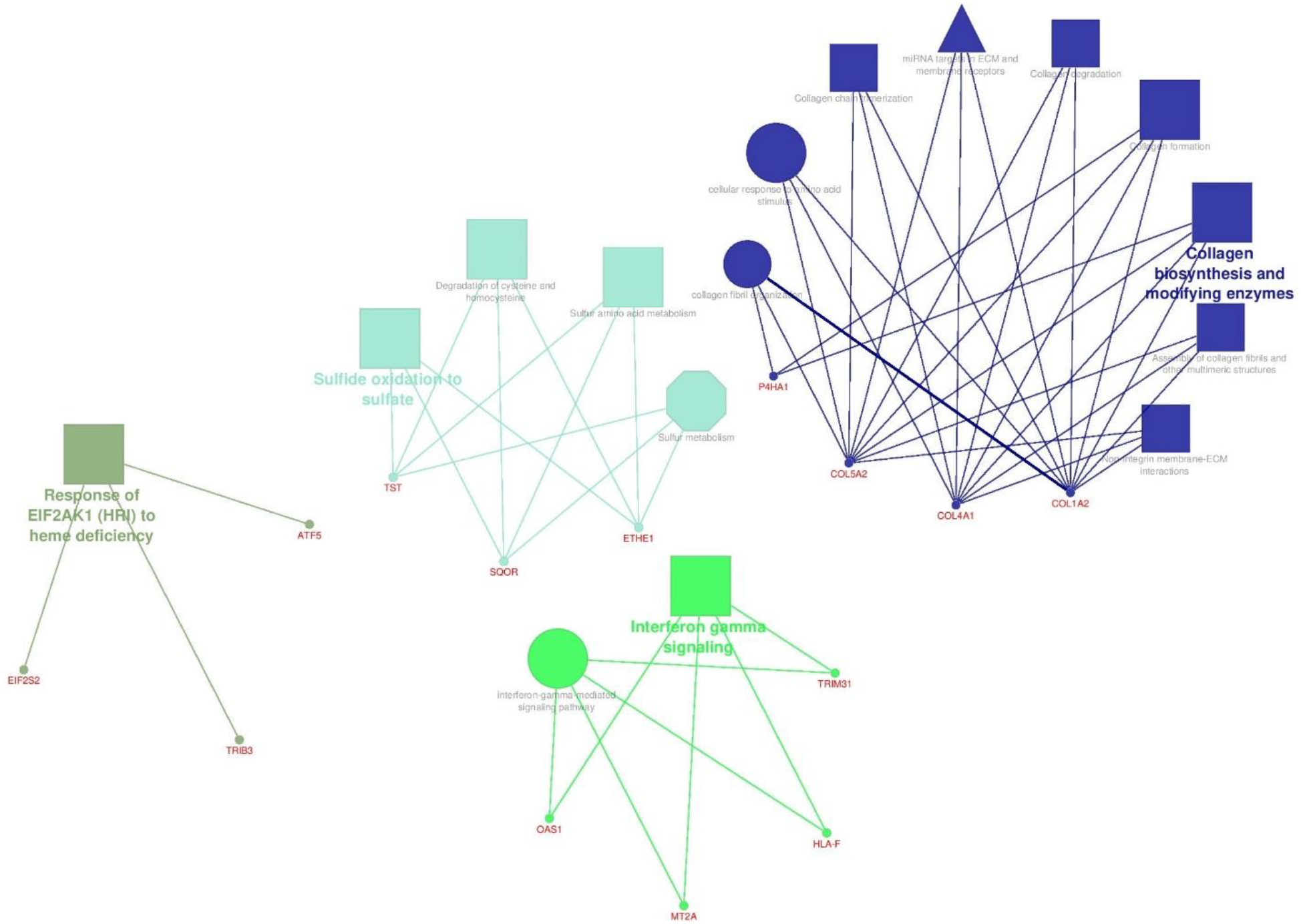
The enrichment results for the core network genes. Terms in the shape of octagons are from KEGG, Triangular terms are from WikiPathways, rectangular expressions are from Reactome and circular terms are from GO database. Size of the terms present their significance.

In the enrichment analysis using “Enrichr” online tool, Gastric Cancer Network2 was of the lowest p-value containing TOP2A and RFC3 genes involved in DNA replication process. Involvement of the same genes in retinoblastoma cancer (WP2446) proposes the potential importance of these genes in different cancers. Top2A was involved in Gastric Cancer Network1 as well. All the enrichment results yielded from “Enrichr” are presented in Supplementary file 5.

### 3.8 Data Integration and Survival Analysis of TCGA Gene Expression Profiles

Expression of DEGs in Table 1 and 2 were further supported by boxplots and survival plots using GEPIA2 web server. Expression profiles were attained from 275 colorectal adenocarcinoma and 41 normal colon RNA-seq samples in TCGA database to create boxplots for each gene in Figure 7. Rather than TWF1, all plots were in agreement with our results. In other words, if a gene was upregulated in our analysis between cancer and normal groups, the expression median for that gene in tumor samples was larger than normal samples in boxplots and vice versa. Even boxplots for expression of some genes that were later shown to be contradictory to other studies, were in favor of our findings. They were MT2A, TRIM31, CDC6, SGK1 and PTP4A1 genes.

**Figure 7:**
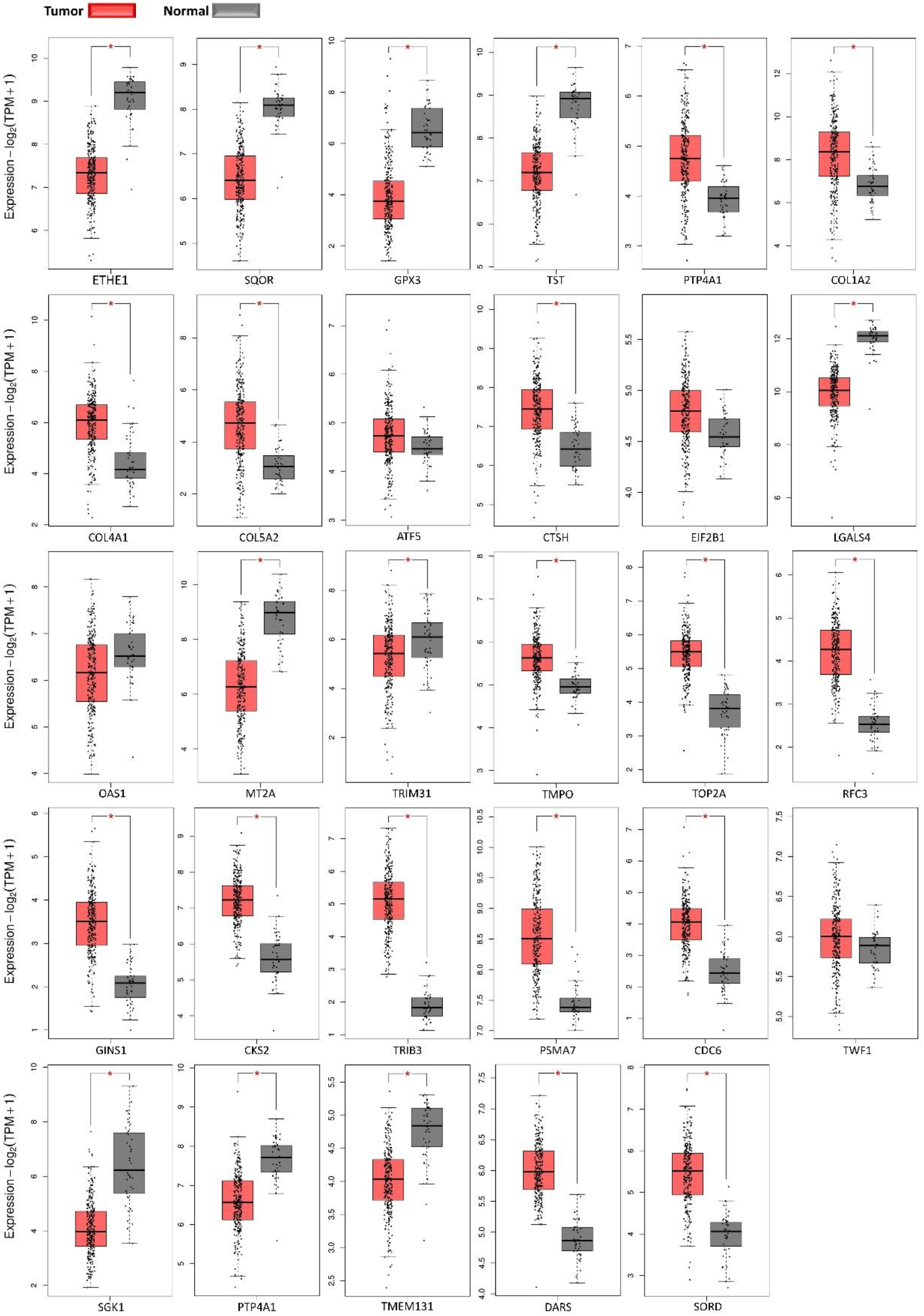
Gene expression profiles boxplots. Red boxes present normalized expression values for tumor adenocarcinoma samples and gray boxes present values for normal colon samples.

Survival plots were also created for DEGs in Table 1 and 2 in different months using TCGA database. Only three genes had a significant p-value larger than 0.05 illustrated in Figure 8 and rest of survival plots were presented in Supplementary file 6. Low expression of LGALS4 is associated with poor survival rate while high expression of COL1A2 and the new reported gene DARS is linked to poor survival rate in colorectal cancer patients. In our study, LGALS4 were downregulated in MvsN and PvsN analyses of all test and validation datasets while DARS and COL1A2 were upregulated in majority of MvsN and PvsN analyses. Although other survival tests were non-significant, majority of them were in agreement with our expression results.

**Figure 8:**
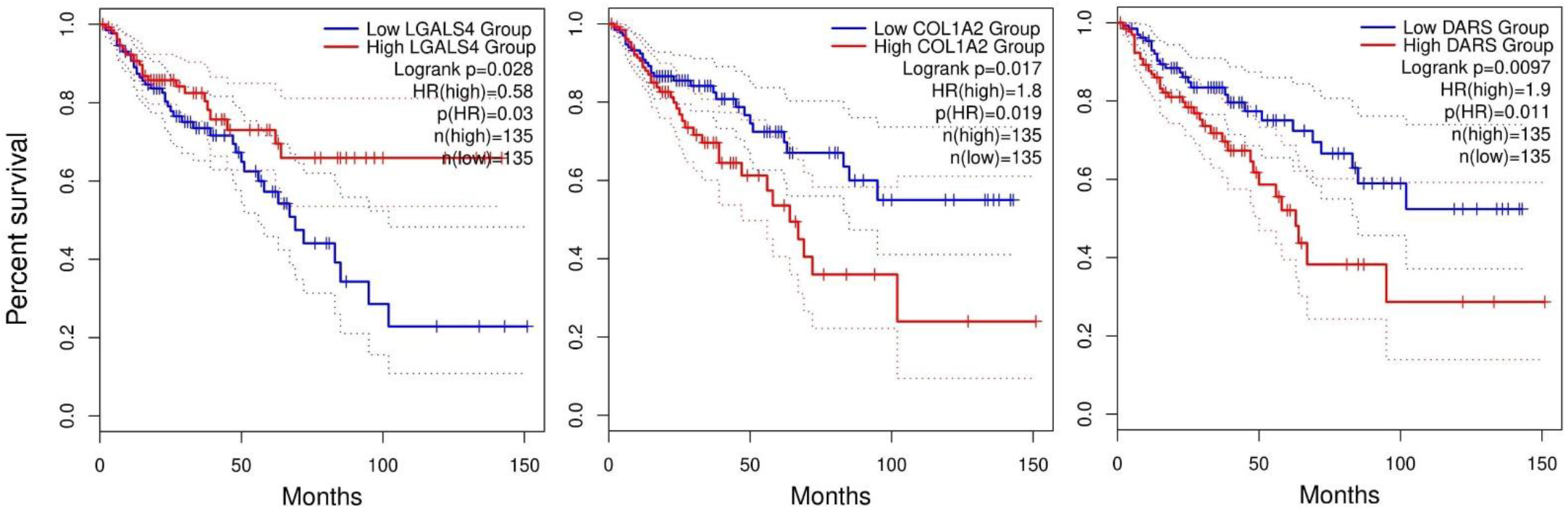
Survival plots. Plots present the monthly survival rate of patients having high expression, red line, or low expression, blue line, of a specific gene. Patients having high expression of LGALS4 had higher survival rates compared to patients having low expression of LGALS4. Contrary, patients having low expression of COL1A2 and DARS genes had higher survival rate

## 4. Discussion

The core network giant component is composed of an up and a down part attached via PLAGL2 transcription factor (TF). The lower part is engaged mainly in cell cycle and DNA replication. Components 2 and 3 contain genes involved in ECM remodeling, component 4 is composed of genes involved in transcription inhibition, and Component 5 is composed of mitochondrial genes playing important roles in controlling cellular redox homeostasis. Here we discussed majority of the genes in the core network exhibiting more similar expression pattern which were present in Table1 and 2.

PLAGL2 is considered as an oncogene in different cancers. It binds to and prevents Pirh2 proteasomal degradation which in turn Pirh2 promotes proteasomal degradation of P53 protein (43). In glioblastoma, PLAGL2 suppresses neural stem cell differentiation by regulating Wnt/β-catenin signaling (44). Besides, PLAGL2 regulates actin cytoskeleton and cell migration through promoting actin stress fibers and focal adhesion (45). Results of PvsN analysis manifests that this gene is induced in primary tumors in colon cancer. In addition this gene had a high betweenness centrality in the giant component (S4). Since this gene connected the two parts of the giant component, it would be a pertinent target for disturbing colon cancer network. Its induction in CRC was supported by the majority of validation datasets in Table 2.

TRIM31 (a ubiquitin ligase) was downregulated in MvsN in all test and validation datasets. However, there are contradictory results in different studies where it was shown to be reduced in lung cancer cells (46) and stepped up in gastric (47) and colorectal cancer (48). Therefore, its downregulation in nine analyses in Tables 1 and 2, needs to be further explored. MT2A gene is an antioxidant which protects cells against hydroxyl radicals and plays an important role in detoxification of heavy metals (49, 50). Expression inhibition of this gene results in proliferation inhibition of CRC cells (51) and its silencing promotes the effects of chemotherapy drugs on osteosarcoma primary tumors (52). However, MT2A gene expression was downregulated in PvsN analyses supported by the results in Table 2. Likewise, it is downregulated in pancreatic cancer as well (53). Therefore, this downregulation in primary CRC tumors has to go under more investigation. OAS1 is a protein induced by interferons which synthesizes the oligomers of adenosine from ATP. These oligomers binds to RNase L to regulate cell growth, differentiation and apoptosis (54). Its expression is downregulated in breast ductal carcinoma and prostate cancer (PCa), at both mRNA and protein levels. In addition, OAS1 expression is negatively correlated with the progression of these cancers (54). The given information supports the downregulation of this gene in our analysis supported by validation datasets. Consequently, expression induction of this gene might help prevent both tumor growth and cell differentiation. The mentioned three genes, TRIM31, MT2A and OAS1, were enriched for IFN-γ and all were downregulated. Although there are contradictory results in different papers, These downregulations at mRNA level would help tumor cells to defeat the anti-cancer properties of interferon gamma signaling.

CTSH gene is a lysosomal cysteine protease upregulated in PvsN. This protease plays an important role in cytoskeletal protein Talin maturation. Talin promotes integrin activation and focal adhesion formation leading to cell migration (55). Upregulation of this gene in CRC was more verified by validation datasets. As a result, suppression of CTSH expression could be a choice of metastasis inhibition. Glutathione peroxidase 3 (GPX3) is an important antioxidant enzyme that protects cells against Reactive Oxygen Species (ROS) and its downregulation occurs in many cancers. For instance, its expression is suppressed in human colon carcinoma Caco2 cell lines, resulting in augmented ROS production (56). It reduces H_2_O_2_ and lipid peroxides to water and lipid alcohols, respectively, and in turn oxidizes glutathione to glutathione disulfide (57). Downregulation of GPX3, happened in PvsN analyses, leading to ascending of H_2_O_2_ level which is positively correlated with tumor progression (58). Its downregulation was further supported by all datasets in Table2. As a result, induction of GPX gene families would be a therapeutic approach.

TMPO gene had the greatest Betweenness centrality illustrating a reduced expression trend in MvsP supported by validation datasets. This gene produces different protein isoforms via alternative splicing (59, 60). The proteins are located in the nucleus of the cells which help form nuclear lamina and maintenance of the nucleus membrane structure (61). TMPO prevents the depolymerization of nuclear laminas and excessive activation of the mitosis pathway. Therefore, its downregulation would prevent excessive mitotic cycle.

TMEM131 is a transmembrane protein that was downregulated in MvsN analyses in all datasets in Table 1 and 2. No documentation was found to connect this gene to cancer. Therefore, this gene would be CRC specific. Furthermore, Enrichment analysis using Enrichr online tool showed that this gene was also involved in interferon gamma signaling (S5).

TOP2A gene was upregulated in PvsN analyses completely endorsed by the validation results. In breast cancer (BC) HER-2 and TOP2A are the molecular targets for several anticancer medicines that are bolstered together (62). Moreover, Copy Number Variations (CNVs) in TOP2A gene have been identified as biomarkers of colorectal cancer (63). This enzyme controls DNA topological structure and its upregulation is a hallmark of aberrant cell growth in tumors (64). TOP2A mRNA expression is an independent prognostic factor in patients with (Estrogen Receptor) ER-positive breast cancer and could be useful in the assessment of breast cancer risk (65). Therefore, in addition to being a possible target for CRC therapy, this gene could be either a possible prognostic or diagnostic marker of CRC.

Replication Factor C subunit 3 (RFC3) plays a role in DNA replication, DNA damage repair, and cell cycle checkpoint control. Hepatocellular carcinoma (HCC) and cell proliferation of ovarian tumors are suppressed by shRNA-mediated silencing of RFC3 gene (66, 67). This gene was upregulated in PvsN analyses and is upregulated in Triple-negative breast cancer (TNBC) as well (68). Its upregulation was more supported by validation datasets. Since expression inhibition of this gene at both mRNA and protein level suppresses the migratory and invasive ability of MCF-7 cell lines (69), this gene would be a therapeutic target for colorectal cancer treatment. Moreover, TOP2A and RFC3 were shown to be engaged in the Gastric Cancer Network2 pathway in the enrichment analysis by Enrichr (S5) which could indicate the importance of these two genes in cancer progression.

Mitotic Arrest Deficient 2 Like1 (MAD2L1) is a mitotic spindle assembly checkpoint molecule upregulated in PvsN in both test and validation analyses. It is responsible for preventing anaphase initiation until precise and complete metaphase alignment of all chromosomes takes place. An increase in the level of MAD2L1 transcripts is detected in a large number of samples with ductal breast carcinoma (70). Its upregulation in our analysis would provide the evidence that cancerous cells were dealing with mitotic deficiencies. The GINS complex is a DNA replication machinery component in the eukaryotes and is an essential tool for initiation and progression of DNA replication forks (71). GINS1 (PSF1) mRNA level is positively correlated with tumor size in CRC patients and is a prognostic marker of CRC (72). This gene has been recently introduced as a targeted oncogenic agent for inhibition of synovial sarcoma (73). It was totally upregulated in PvsN analyses in both Tables 1 and 2. therefore, its expression inhibition would be a potential target for inhibition of tumor growth by disturbing DNA replication machinery.

CDC6, one of the core genes, plays a critical role in regulation of the eukaryotic DNA replication onset and its downregulation has been demonstrated in prostate cancer (74). It is a regulator of cell cycle in S phase and its expression is regulated by E2F and androgen receptor (AR) in PCa cells (75). Transfection of CDC6 siRNA leads to not only decreased level of ovarian cancer cell proliferation but also increased apoptosis rates (76). Cdc6 along with Cdt1 are highly expressed in aggressive BC and therefore is considered as a potent therapeutic target in BC patients (77). Results for this gene in MvsP were totally contradictory to the BC results but is similar to prostate cancer. Majority of validation datasets depicted downregulation of this gene in CRC. No study directly measured the expression level of this gene in CRC samples therefore it is worth investigation to see whether it could be a CRC biomarker or a curative target.

CKS2 protein interacts with the catalytic subunit of the cyclin-dependent kinases and its downregulation contributes to suppression of p-Akt and p-mTOR. Therefore, one of CSK2 oncogenic roles is played by Akt/mTOR oncogenic pathway (78). CKS2 is expressed at high level in CRC tissues and it has revealed that increased CKS2 expression is highly correlated with enhanced metastatic stage (79). Importantly, CKS2 is considered as a potential biomarker and therapeutic target for the BC treatment due to the fact that its inhibition suppresses cell proliferation and invasion in vitro and in vivo (80). In the PvN analysis, this gene was upregulated which would be a therapeutic target for CRC treatment because validation results completely supported this upregulation.

PSMA7 gene encodes a protein that is one of the essential subunits of 20S proteasome complex (81). Overexpression of PSMA7 both at the mRNA and protein levels has been reported in gastric cancer (82). Depletion of PSMA7 by shRNA-transfected RKO CRC cell lines mediates inhibition of cell growth and migration. Consequently, inhibition of PSMA7 could be a beneficial therapeutic strategy for colorectal cancer patients (83). This gene was upregulated in PvsN analyses in test and validation datasets.

DARS was found to be upregulated in MvsN and PvsN analyses in all test and validation datasets (total of 16 analyses). This gene encodes a member of a multi-enzyme complex that its role has been proved in mediating attachment of amino acids to their cognate tRNAs. No study has investigated the role of this gene in cancer progression so far. Only two studies have reported that DARS-AS1 gene is positively associated with the pathological stages in thyroid and ovarian cancer by targeting mir-129 and mir-532-3p respectively (84, 85). This gene might be a CRC prognostic marker or a curative target.

EIF-2 consists of alpha, beta, and gamma subunits. EIF2B or EIF2S2 acts in the early steps of protein synthesis. GTP-bound EIF-2 transfers Met-tRNAi to the 40S ribosomal subunit to start protein synthesis. The hydrolysis of GTP to GDP takes place at the end of the initiation process that leads to release of the inactive eIF2·GDP from ribosome. Exchange of GDP for GTP is performed by beta subunit so that active EIF-2 is ready for another round of initiation (86). In one study, EIF2B was proposed as a potent therapeutic target in lung cancer [76]. Moreover, elimination of natural killer cell cytotoxicity via promoted expression of natural killer (NK) cell ligands is done by pSer535-eIF2B following the expression of pSer9-GSK-3β (inactive GSK3β) and generation of ROS which promotes breast cancer growth and metastasis (87). Since Tyr216-GSK-3β (Active GSK3β) has the inhibitory effects on the EMT process by interfering with TNF-alpha signaling (88), induction of active GSK-3β together with suppression of EIF2B would be a therapeutic approach to prevent EMT (89). EIF2B stepped up in PvsN analyses which was supported by validation results.

TWF1 gene encodes twinfilin, an actin monomer-binding protein that promotes EMT in pancreatic cancer tissues (90). siRNAs of TWF1 dramatically inhibits F-actin organization and focal adhesions formation that promotes the mesenchymal-to-epithelial transition (MET) in MDA-MB-231 cell lines. In addition, The responsiveness of these cell lines to anti-cancer drugs such as doxorubicin and paclitaxel is amplified by inhibition of TWF1 expression by both microRNA and siRNA (91). Furthermore, expression levels of EMT markers, VIM and SNAI2, are reduced as a result of miR-30c action on TWF1 mRNA (91). However, in MvsN analyses, this gene witnessed a decreased expression in both test and validation datasets. As a results, Its upregulation in CRC has to be further explored.

SGK1, a member of component 2, and AKT are two families of AGC protein superfamily. SGK1 is a serine/threonine kinase that activates certain potassium, sodium, and chloride channels (92). SGK1 is a downstream effector of PI3K, which runs pathways independent of pathways shared with AKT. The two kinases are phosphorylated and activated by PDK1 and mTORC2 complex (93, 94). In general, PI3K-dependent survival signals can be mediated by either Akt or SGK1 that inactivates the pro-apoptotic proteins Bad and FKHRL1 (95). In a study on A498 kidney cancer cells, they found that survival signals promoted by IL-2 is mediated by SGK1 activation (96). Moreover, the promoter of SGK1 is under tight control of the p53 protein (97). SGK1 has been shown to mediate cell survival and drug resistance to platinoid and taxane compounds in breast cancer patients (98). On the contrary, This gene was totally downregulated in PvsN analyses in all validation and test datasets. These overall downregulations might be specific to CRC so it could be a diagnostic hallmark of CRC and should go under more interrogation.

Component 3 contains collagen (COL1A2, COL5A2 and COL4A1) and P4HA1 (a collagen hydroxylase) genes interconnected in the process of ECM remodeling based on the enrichment results. All members witnessed an ascending trend in expression from normal samples to metastatic samples in Figure 5 panels. In test datasets, collagen genes presented an upregulation trend in MvsN and PvsN analyses, while their expression followed a mixed trend in validation datasets. P4HA1 one of the core genes upregulated in MvsP in all test and validation datasets. Only expression of COL1A2 followed a homogeneous upregulating trend in both test and validation datasets which is a marker of EMT as well (99). P4HA1 is engaged in breast and pancreatic metastasis (100, 101). Under the tumor hypoxic conditions, HIF-1 induces expression of genes that encodes collagen prolyl (P4HA1 and P4HA2) and lysyl (PLOD2) hydroxylases. P4HA1 and P4HA2 are required for collagen deposition, whereas PLOD2 is required for ECM stiffening and collagen fiber alignment (102). These changes in ECM triggered by HIF-1 are necessary for motility and invasion because in focal adhesion junctions, actin cytoskeleton is connected to ECM through attachment of integrins to collagens (103). Besides, there is a positive feedback between P4HA1 and HIF-1 in modulation of ECM. As a result, targeting P4HA1 and P4HA2 expressions would inhibit the progression of cell migration via HIF1-Collagen pathway.

PTP4A1 a member of component 4, is a protein phosphatase engaged in p21-activated kinase (PAK) signaling pathway. Inhibition of PTP4A1 gene in MDA-MB-231 breast cancer cell lines by an increase in miR-944 expression impairs cell invasion (104). However, This gene was downregulated in MvsN and PvsN in all test datasets and in the majority of validation datasets. This downregulation would be a biomarker for CRC and its molecular role in CRC needs to be interrogated. BCL-2 is a target of ATF5 one of the core genes (105). ATF5 was upregulated in MvsP analyses in both test and validation datasets. There are pieces of evidence that link the role of ATF5 in mitochondrial dysfunction in cancer progression (106). In malignant glioma, metastatic cells take advantage of survival signals triggered by ATF5 gene which is important to ignore anchorage-dependent and niche-dependent cell death signals (107). As a results, expression inhibition of ATF5 would hinder the survival signals in CRC cells. TRIB3 is a prognosis hallmark of colorectal cancer activated under hypoxic conditions (108). TRIB3 silencing suppresses VEGF-A expression in gastric cancerous cells which inhibits endothelial cell migration and vessel formation. This gene was upregulated in MvsN analyses in all test and validation datasets, therefore, it would be a favorable target for anti-angiogenic therapy (109).

Genes in component 5 are mitochondrial which their role in cancer progression has not been sufficiently investigated so far. All three genes were downregulated in our analysis in both validation and test datasets. They also exhibited totally a reducing trend from normal to primary and from primary to metastatic in Figure 5 panels. These genes are highly expressed in normal colon tissue compared to other tissues due to the presence of anaerobic bacteria in the digestive tract (110). These findings are supported by the RNA-seq expression information in the Gene database of NCBI (111). ETHE1 (persulfide dioxygenase) and SQOR are antioxidants that convert hydrogen sulfide (H2S) at first to persulfide then to sulfite. Therefore, they protect cells against toxic concentrations of sulfide. ETHE1 gene was downregulated in the three analyses while SQOR was downregulated in MvsN and PvsN analyses. All these expressions were totally verified by the validation datasets. Their downregulation is essential for cancer cells proliferation and survival since under the hypoxic environment of CRC tumor, sulfide is a supplementary tool that provides electron for mitochondrial electron transport chain (ETC) to generate ATP (112). These mechanisms along with Warburg effect help tumor cells to survive the hypoxic environment. As a result, helping expression induction or activation of ETHE1 and SQOR proteins will increase sulfide scavenging and this would hinder CRC tumor growth. TST thiosulfate sulfurtransferase encodes a protein that is localized to the mitochondria and catalyzes the conversion of thiosulfate and cyanide to thiocyanate and sulfite respectively. Therefore, like the previous two mitochondrial enzymes, it acts in Hydrogen sulfide metabolism (113).

SORD is another element of component 6 upregulated in MvsN and PvsN analyses. The connection of this gene with cancer has not been efficiently investigated. Since it exhibited a totally ascending trend in all validation and test datasets and in Figure 5, it might be a potential target and biomarker for CRC treatment. LGALS4 is implicated in regulating cell-cell and cell-matrix interactions so its induction might have positive curative impacts on CRC cells. This gene is mostly expressed in small intestine, colon, and rectum, which is suppressed in CRC (114). It was downregulated in MvsN and PvsN analyses in both validation and test datasets. It is also a blood marker of CRC (115).

In summary, we illustrated some therapeutic targets and biomarkers for CRC. A combination of these targets would beneficially disturb progression of colorectal cancer. Generally, all the discovered antioxidants were downregulated in different stages of CRC namely ETHE1, SQOR, TST and GPX3. We proposed that these downregulations under hypoxic conditions would help cancer cells to produce more energy for cell proliferation. In addition, the hypoxic environment alters ECM suitable for cell migration by induction of P4HA1 gene through HIF-1 signaling pathway and induction of COL1A2. Boxplots (expression profiling) in figure 7 supported our results for all these genes. In addition, survival plot in Figure 8 demonstrated that there is a higher death probability for CRC patients expressing high level of COL1A2 than patients having low level of this gene. Consequently, colorectal cancer cells would take advantages of explained mechanisms along with Warburg effect to not only survive from the hypoxic environment of tumors but also proliferate faster and migrate better. Therefore, induction of mentioned anti-oxidants and suppression of P4HA1 and COL1A2 genes would be a choice of CRC treatment.

Induction of active GSK-3β together with suppression of EIF2B would prevent EMT in CRC. Induction of OAS1 to increase the anti-cancer effects of interferon gamma, suppression of CTSH to hinder formation of focal adhesions, expression inhibition of ATF5 gene to make cancer cells sensitive to anchorage-dependent death signals and induction of LGALS4 gene (supported by survival analysis) to recover cell-cell junctions would be the combination of genetic targets that prevent EMT and cell migration. In addition, expression inhibition of TMPO, TOP2A, RFC3, GINS1 and CKS2 genes could prevent tumor growth and TRIB3 expression suppression would be a favorable target for anti-angiogenic therapy. PSMA7 gene was a previously reported target for CRC treatment that was also found in our study. Results for expression of all these genes were supported by expression profiling.

MT2A and TRIM31 which were engaged in IFN-γ signaling, CDC6, SGK1 and PTP4A1 genes presented a homogeneous expression pattern in both test and validation datasets although our results were contradictory to other studies in different cancers. Nevertheless, we used 10 different datasets from different technologies to ensure the accuracy of the results. Besides, expression profiling totally supported expression of these genes. They have to be further interrogated in colorectal cancer progression.

TMEM131, DARS and SORD genes had specific uniform expression trends as analyses went from normal to metastatic. DARS expression inhibition would increase the survival rate in CRC patients based on Figure 8. The relation of these three genes to colorectal cancer progression has been reported for the first time in this study, hence more investigation is required to find their molecular mechanism causing colorectal cancer promotion.

## References

1. Favoriti P, Carbone G, Greco M, Pirozzi F, Pirozzi REM, Corcione F. Worldwide burden of colorectal cancer: a review. Updates in surgery. 2016;68(1):7–11.

2. Ferlay J, Shin H, Bray F, Forman D, Mathers C, Parkin DG. v2. 0, Cancer Incidence and Mortality Worldwide. IARC Cancerbase No. 2008;10.

3. Chambers A, Groom A, MacDonald I. ā€ Dissemination and Growth of Cancer Cells in Metastatic Sites, ā€ Nat. Rev Cancer. 2002;2:563ā.

4. Jamora C, Fuchs E. Intercellular adhesion, signalling and the cytoskeleton. Nature cell biology. 2002;4(4):E101.

5. Al-Sohaily S, Biankin A, Leong R, Kohonen-Corish M, Warusavitarne J. Molecular pathways in colorectal cancer. Journal of gastroenterology and hepatology. 2012;27(9):1423–31.

6. Sierra JR, Cepero V, Giordano S. Molecular mechanisms of acquired resistance to tyrosine kinase targeted therapy. Molecular cancer. 2010;9(1):75.

7. Wang S-W, Sun Y-M. The IL-6/JAK/STAT3 pathway: potential therapeutic strategies in treating colorectal cancer. International journal of oncology. 2014;44(4):1032–40.

8. Dihlmann S, von Knebel Doeberitz M. Wnt/β-catenin-pathway as a molecular target for future anti-cancer therapeutics. International journal of cancer. 2005;113(4):515–24.

9. Krüger A, Soeltl R, Sopov I, Kopitz C, Arlt M, Magdolen V, et al. Hydroxamate-type matrix metalloproteinase inhibitor batimastat promotes liver metastasis. Cancer research. 2001;61(4):1272–5.

10. Katz LH, Li Y, Chen J-S, Muñoz NM, Majumdar A, Chen J, et al. Targeting TGF-β signaling in cancer. Expert opinion on therapeutic targets. 2013;17(7):743–60.

11. Pickup M, Novitskiy S, Moses HL. The roles of TGFβ in the tumour microenvironment. Nature Reviews Cancer. 2013;13(11):788–99.

12. Rychahou P, Bae Y, Reichel D, Zaytseva YY, Lee EY, Napier D, et al. Colorectal cancer lung metastasis treatment with polymer—drug nanoparticles. Journal of controlled release. 2018;275:85–91.

13. Rychahou P, Bae Y, Zaytseva Y, Lee EY, Weiss HL, Evers BM. Colorectal cancer pulmonary metastasis treatment with lung-selective delivery of PAN-class I PI3K inhibitors. AACR; 2016.

14. Jonker DJ, O’Callaghan CJ, Karapetis CS, Zalcberg JR, Tu D, Au H-J, et al. Cetuximab for the treatment of colorectal cancer. New England Journal of Medicine. 2007;357(20):2040–8.

15. Tol J, Koopman M, Cats A, Rodenburg CJ, Creemers GJ, Schrama JG, et al. Chemotherapy, bevacizumab, and cetuximab in metastatic colorectal cancer. New England Journal of Medicine. 2009;360(6):563–72.

16. Gautier L, Cope L, Bolstad BM, Irizarry RA. affy–analysis of Affymetrix GeneChip data at the probe level. Bioinformatics. 2004;20(3):307–15.

17. Do JH, Choi D-K. Normalization of microarray data: single-labeled and dual-labeled arrays. Molecules & Cells (Springer Science & Business Media BV). 2006;22(3).

18. Ramasamy A, Mondry A, Holmes CC, Altman DG. Key issues in conducting a meta-analysis of gene expression microarray datasets. PLoS medicine. 2008;5(9):e184.

19. Gentleman R, Carey V, Huber W, Hahne F. Genefilter: Methods for filtering genes from microarray experiments. R package version. 2011;1(0).

20. Smyth GK. Limma: linear models for microarray data. Bioinformatics and computational biology solutions using R and Bioconductor: Springer; 2005 p. 397–420.

21. Ferreira J, Zwinderman A. On the Benjamini—Hochberg method. The Annals of Statistics. 2006;34(4):1827–49.

22. Love MI, Huber W, Anders S. Moderated estimation of fold change and dispersion for RNA-seq data with DESeq2. Genome biology. 2014;15(12):550.

23. Luke DA. A user’s guide to network analysis in R: Springer; 2015.

24. Langfelder P, Horvath S. WGCNA: an R package for weighted correlation network analysis. BMC bioinformatics. 2008;9(1):559.

25. Jasika N, Alispahic N, Elma A, Ilvana K, Elma L, Nosovic N, editors. Dijkstra’s shortest path algorithm serial and parallel execution performance analysis. 2012 proceedings of the 35th international convention MIPRO; 2012: IEEE.

26. Berger SI, Ma’ayan A, Iyengar R. Systems pharmacology of arrhythmias. Sci Signal. 2010;3(118):ra30–ra.

27. Azimzadeh Jamalkandi S, Mozhgani S-H, Gholami Pourbadie H, Mirzaie M, Noorbakhsh F, Vaziri B, et al. Systems biomedicine of rabies delineates the affected signaling pathways. Frontiers in microbiology. 2016;7:1688.

28. Bindea G, Mlecnik B, Hackl H, Charoentong P, Tosolini M, Kirilovsky A, et al. ClueGO: a Cytoscape plug-in to decipher functionally grouped gene ontology and pathway annotation networks. Bioinformatics. 2009;25(8):1091–3.

29. Kanehisa M, Goto S. KEGG: kyoto encyclopedia of genes and genomes. Nucleic acids research. 2000;28(1):27–30.

30. Croft D, O’Kelly G, Wu G, Haw R, Gillespie M, Matthews L, et al. Reactome: a database of reactions, pathways and biological processes. Nucleic acids research. 2010;39(suppl_1):D691–D7.

31. Pico AR, Kelder T, Van Iersel MP, Hanspers K, Conklin BR, Evelo C. WikiPathways: pathway editing for the people. PLoS Biol. 2008;6(7):e184.

32. Kuleshov MV, Jones MR, Rouillard AD, Fernandez NF, Duan Q, Wang Z, et al. Enrichr: a comprehensive gene set enrichment analysis web server 2016 update. Nucleic acids research. 2016;44(W1):W90–W7.

33. Tang Z, Kang B, Li C, Chen T, Zhang Z. GEPIA2: an enhanced web server for large-scale expression profiling and interactive analysis. Nucleic acids research. 2019;47(W1):W556–W60.

34. Weinstein JN, Collisson EA, Mills GB, Shaw KRM, Ozenberger BA, Ellrott K, et al. The cancer genome atlas pan-cancer analysis project. Nature genetics. 2013;45(10):1113.

35. Oldham MC, Horvath S, Konopka G, Iwamoto K, Langfelder P, Kato T, et al. Identification and Removal of Outlier Samples Supplement for:“ Functional Organization of the Transcriptome in Human Brain. dim (dat1).1(18631):105.

36. Frantz C, Stewart KM, Weaver VM. The extracellular matrix at a glance. Journal of cell science. 2010;123(24):4195–200.

37. Huang S, Ingber DE. Cell tension, matrix mechanics, and cancer development. Cancer cell. 2005;8(3):175–6.

38. Gooch JL, Herrera RE, Yee D. The role of p21 in interferon gamma-mediated growth inhibition of human breast cancer cells. Cell growth & differentiation: the molecular biology journal of the American Association for Cancer Research. 2000;11(6):335.

39. Li P, Du Q, Cao Z, Guo Z, Evankovich J, Yan W, et al. Interferon-gamma induces autophagy with growth inhibition and cell death in human hepatocellular carcinoma (HCC) cells through interferon-regulatory factor-1 (IRF-1). Cancer letters. 2012;314(2):213–22.

40. Wang L, Wang Y, Song Z, Chu J, Qu X. Deficiency of interferon-gamma or its receptor promotes colorectal cancer development. Journal of Interferon & Cytokine Research. 2015;35(4):273–80.

41. Zaidi MR, Merlino G. The two faces of interferon-γ in cancer. Clinical cancer research. 2011;17(19):6118–24.

42. Mojic M, Takeda K, Hayakawa Y. The dark side of IFN-γ: its role in promoting cancer immunoevasion. International journal of molecular sciences. 2018;19(1):89.

43. Zheng G, Ning J, Yang Y-C. PLAGL2 controls the stability of Pirh2, an E3 ubiquitin ligase for p53. Biochemical and biophysical research communications. 2007;364(2):344–50.

44. Zheng H, Ying H, Wiedemeyer R, Yan H, Quayle SN, Ivanova EV, et al. PLAGL2 regulates Wnt signaling to impede differentiation in neural stem cells and gliomas. Cancer cell. 2010;17(5):497–509.

45. Sekiya R, Maeda M, Yuan H, Asano E, Hyodo T, Hasegawa H, et al. PLAGL2 regulates actin cytoskeletal architecture and cell migration. Carcinogenesis. 2014;35(9):1993–2001.

46. Li H, Zhang Y, Zhang Y, Bai X, Peng Y, He P. TRIM31 is downregulated in non-small cell lung cancer and serves as a potential tumor suppressor. Tumor Biology. 2014;35(6):5747–52.

47. Sugiura T. The cellular level of TRIM31, an RBCC protein overexpressed in gastric cancer, is regulated by multiple mechanisms including the ubiquitin–proteasome system. Cell biology international. 2011;35(7):657–61.

48. Wang H, Yao L, Gong Y, Zhang B. TRIM31 regulates chronic inflammation via NF-κB signal pathway to promote invasion and metastasis in colorectal cancer. American journal of translational research. 2018;10(4):1247.

49. Ma H, Su L, Yue H, Yin X, Zhao J, Zhang S, et al. HMBOX1 interacts with MT2A to regulate autophagy and apoptosis in vascular endothelial cells. Scientific reports. 2015;5:15121.

50. Liu Y, Liu H, Chen W, Yang T, Zhang W. EOLA1 protects lipopolysaccharide induced IL-6 production and apoptosis by regulation of MT2A in human umbilical vein endothelial cells. Molecular and cellular biochemistry. 2014;395(1-2):45–51.

51. Xu Y, Jin H, Tan X, Liu X, Ding Y. Tea polyphenol inhibits colorectal cancer with microsatellite instability by regulating the expressions of HES1, JAG1, MT2A and MAFA. Zhong xi yi jie he xue bao= Journal of Chinese integrative medicine. 2010;8(9):870–6.

52. Mangelinck A, da Costa MEM, Stefanovska B, Bawa O, Polrot M, Gaspar N, et al. MT2A is an early predictive biomarker of response to chemotherapy and a potential therapeutic target in osteosarcoma. Scientific reports. 2019;9(1):1–12.

53. Hui B, Xu Y, Zhao B, Ji H, Ma Z, Xu S, et al. Overexpressed long noncoding RNA TUG1 affects the cell cycle, proliferation, and apoptosis of pancreatic cancer partly through suppressing RND3 and MT2A. OncoTargets and therapy. 2019;12:1043.

54. Maia CJ, Rocha SM, Socorro S, Schmitt F, Santos CR. Oligoadenylate synthetase 1 (OAS1) expression in human breast and prostate cancer cases, and its regulation by sex steroid hormones. Advances in Modern Oncology Research. 2016;2(2):97–110.

55. Jevnikar Z, Rojnik M, Jamnik P, Doljak B, Fonović UP, Kos J. Cathepsin H mediates the processing of talin and regulates migration of prostate cancer cells. Journal of Biological Chemistry. 2013;288(4):2201–9.

56. Barrett CW, Ning W, Chen X, Smith JJ, Washington MK, Hill KE, et al. Tumor suppressor function of the plasma glutathione peroxidase gpx3 in colitis-associated carcinoma. Cancer research. 2013;73(3):1245–55.

57. Sarsour EH, Kumar MG, Chaudhuri L, Kalen AL, Goswami PC. Redox control of the cell cycle in health and disease. Antioxidants & redox signaling. 2009;11(12):2985–3011.

58. Moloney JN, Cotter TG, editors. ROS signalling in the biology of cancer. Seminars in cell & developmental biology; 2018: Elsevier.

59. Holmer L, Worman H. Inner nuclear membrane proteins: functions and targeting. Cellular and Molecular Life Sciences CMLS. 2001;58(12-13):1741–7.

60. Harris CA, Andryuk PJ, Cline S, Chan HK, Natarajan A, Siekierka JJ, et al. Three distinct human thymopoietins are derived from alternatively spliced mRNAs. Proceedings of the National Academy of Sciences. 1994;91(14):6283–7.

61. Huang W, Su X, Yan W, Kong Z, Wang D, Huang Y, et al. Overexpression of AR-regulated lncRNA TMPO-AS1 correlates with tumor progression and poor prognosis in prostate cancer. The Prostate. 2018;78(16):1248–61.

62. Simon R, Atefy R, Wagner U, Forster T, Fijan A, Bruderer J, et al. HER-2 and TOP2A coamplification in urinary bladder cancer. International journal of cancer. 2003;107(5):764–72.

63. Vang Nielsen K, Ejlertsen B, Møller S, Trøst Jørgensen J, Knoop A, Knudsen H, et al. The value of TOP2A gene copy number variation as a biomarker in breast cancer: Update of DBCG trial 89D. Acta oncologica. 2008;47(4):725–34.

64. De Resende MF, Vieira S, Chinen LTD, Chiappelli F, da Fonseca FP, Guimarães GC, et al. Prognostication of prostate cancer based on TOP2A protein and gene assessment: TOP2A in prostate cancer. Journal of translational medicine. 2013;11(1):36.

65. Rody A, Karn T, Ruckhäberle E, Müller V, Gehrmann M, Solbach C, et al. Gene expression of topoisomerase II alpha (TOP2A) by microarray analysis is highly prognostic in estrogen receptor (ER) positive breast cancer. Breast cancer research and treatment. 2009;113(3):457–66.

66. Shen H, Xu J, Zhao S, Shi H, Yao S, Jiang N. ShRNA-mediated silencing of the RFC3 gene suppress ovarian tumor cells proliferation. International journal of clinical and experimental pathology. 2015;8(8):8968.

67. Yao Z, Hu K, Huang H, Xu S, Wang Q, Zhang P, et al. shRNA-mediated silencing of the RFC3 gene suppresses hepatocellular carcinoma cell proliferation. International journal of molecular medicine. 2015;36(5):1393–9.

68. He Z-Y, Wu S-G, Peng F, Zhang Q, Luo Y, Chen M, et al. Up-Regulation of RFC3 promotes triple negative breast cancer metastasis and is associated with poor prognosis via EMT. Translational oncology. 2017;10(1):1–9.

69. Zhou J, Zhang W-W, Peng F, Sun J-Y, He Z-Y, Wu S-G. Downregulation of hsa_circ_0011946 suppresses the migration and invasion of the breast cancer cell line MCF-7 by targeting RFC3. Cancer management and research. 2018;10:535.

70. Scintu M, Vitale R, Prencipe M, Gallo AP, Bonghi L, Valori VM, et al. Genomic instability and increased expression of BUB1B and MAD2L1 genes in ductal breast carcinoma. Cancer letters. 2007;254(2):298–307.

71. Labib K, Gambus A. A key role for the GINS complex at DNA replication forks. Trends in cell biology. 2007;17(6):271–8.

72. Wei H-B, Wen J-Z, Wei B, Han X-Y, Zhang S. Expression and clinical significance of GINS complex in colorectal cancer. Zhonghua wei chang wai ke za zhi= Chinese journal of gastrointestinal surgery. 2011;14(6):443–7.

73. Tang L, Yu W, Wang Y, Li H, Shen Z. Anlotinib inhibits synovial sarcoma by targeting GINS1: a novel downstream target oncogene in progression of synovial sarcoma. Clinical and Translational Oncology. 2019:1–10.

74. Robles LD, Frost AR, Davila M, Hutson AD, Grizzle WE, Chakrabarti R. Down-regulation of Cdc6, a cell cycle regulatory gene, in prostate cancer. Journal of Biological Chemistry. 2002;277(28):25431–8.

75. Liu Y, Gong Z, Sun L, Li X. FOXM1 and androgen receptor co-regulate CDC6 gene transcription and DNA replication in prostate cancer cells. Biochimica et Biophysica Acta (BBA)-Gene Regulatory Mechanisms. 2014;1839(4):297–305.

76. Sun T-Y, Xie H-J, Li Z, He H, Kong L-F. Expression of CDC6 in ovarian cancer and its effect on proliferation of ovarian cancer cells. Int J Clin Exp Med. 2016;9(6):10544–50.

77. Mahadevappa R, Neves H, Yuen SM, Bai Y, McCrudden CM, Yuen HF, et al. The prognostic significance of Cdc6 and Cdt1 in breast cancer. Scientific reports. 2017;7(1):985.

78. Xu J, Wang Y, Xu D. CKS2 promotes tumor progression and metastasis and is an independent predictor of poor prognosis in epithelial ovarian cancer. European review for medical and pharmacological sciences. 2019;23(8):3225–34.

79. Yu M-H, Luo Y, Qin S-L, Wang Z-S, Mu Y-F, Zhong M. Up-regulated CKS2 promotes tumor progression and predicts a poor prognosis in human colorectal cancer. American journal of cancer research. 2015;5(9):2708.

80. Huang N, Wu Z, Hong H, Wang X, Yang F, Li H. Overexpression of CKS2 is associated with a poor prognosis and promotes cell proliferation and invasion in breast cancer. Molecular medicine reports. 2019;19(6):4761–9.

81. Coux O, Tanaka K, Goldberg ALJArob. Structure and functions of the 20S and 26S proteasomes. 1996;65(1):801–47.

82. Xia S, Tang Q, Wang X, Zhang L, Jia L, Wu D, et al. Overexpression of PSMA7 predicts poor prognosis in patients with gastric cancer. Oncology letters. 2019;18(5):5341–9.

83. Hu X-T, Chen W, Zhang F-B, Shi Q-L, Hu J-B, Geng S-M, et al. Depletion of the proteasome subunit PSMA7 inhibits colorectal cancer cell tumorigenicity and migration. Oncology reports. 2009;22(5):1247–52.

84. Zheng W, Tian X, Cai L, Shen Y, Cao Q, Yang J, et al. LncRNA DARS-AS1 regulates microRNA-129 to promote malignant progression of thyroid cancer. European Review for Medical and Pharmacological Sciences. 2019;23:10443–52.

85. Huang K, Fan W, Fu X, Li Y, Meng Y. Long noncoding RNA DARS-AS1 acts as an oncogene by targeting miR-532-3p in ovarian cancer. European review for medical and pharmacological sciences. 2019;23(6):2353–9.

86. Kimball SR. Eukaryotic initiation factor eIF2. The international journal of biochemistry & cell biology. 1999;31(1):25–9.

87. Jin F, Wu Z, Hu X, Zhang J, Gao Z, Han X, et al. The PI3K/Akt/GSK-3β/ROS/eIF2B pathway promotes breast cancer growth and metastasis via suppression of NK cell cytotoxicity and tumor cell susceptibility. Cancer biology & medicine. 2019;16(1):38.

88. Wang H, Wang H-S, Zhou B-H, Li C-L, Zhang F, Wang X-F, et al. Epithelial—mesenchymal transition (EMT) induced by TNF-α requires AKT/GSK-3β-mediated stabilization of snail in colorectal cancer. PloS one. 2013;8(2):e56664.

89. Goto D, Tanaka I, Sato M, Kato T, Miyazawa A, Hase T, et al. P1.03-13 eIF2β, A Subunit of Translation-Initiation Factor EIF2, as a Potential Therapeutic Target for Non-Small Cell Lung Cancer. Journal of Thoracic Oncology. 2018;13(10):S516–S7.

90. Jin X, Pang W, Zhang Q, Huang H. MicroRNA-486-5p improves nonsmall-cell lung cancer chemotherapy sensitivity and inhibits epithelial—mesenchymal transition by targeting twinfilin actin binding protein 1. Journal of International Medical Research. 2019:0300060519850739.

91. Bockhorn J, Dalton R, Nwachukwu C, Huang S, Prat A, Yee K, et al. MicroRNA-30c inhibits human breast tumour chemotherapy resistance by regulating TWF1 and IL-11. Nature communications. 2013;4:1393.

92. Arencibia JM, Pastor-Flores D, Bauer AF, Schulze JO, Biondi RM. AGC protein kinases: from structural mechanism of regulation to allosteric drug development for the treatment of human diseases. Biochimica Et Biophysica Acta (BBA)-Proteins and Proteomics. 2013;1834(7):1302–21.

93. Di Cristofano A. SGK1: the dark side of PI3K signaling. Current topics in developmental biology. 123: Elsevier; 2017 p. 49–71.

94. Park J, Leong ML, Buse P, Maiyar AC, Firestone GL, Hemmings BA. Serum and glucocorticoid-inducible kinase (SGK) is a target of the PI 3-kinase-stimulated signaling pathway. The EMBO journal. 1999;18(11):3024–33.

95. Brunet A, Park J, Tran H, Hu LS, Hemmings BA, Greenberg ME. Protein kinase SGK mediates survival signals by phosphorylating the forkhead transcription factor FKHRL1 (FOXO3a). Molecular and cellular biology. 2001;21(3):952–65.

96. Amato R, Menniti M, Agosti V, Boito R, Costa N, Bond HM, et al. IL-2 signals through Sgk1 and inhibits proliferation and apoptosis in kidney cancer cells. Journal of molecular medicine. 2007;85(7):707–21.

97. Maiyar AC, Huang AJ, Phu PT, Cha HH, Firestone GL. p53 stimulates promoter activity of the sgk serum/glucocorticoid-inducible serine/threonine protein kinase gene in rodent mammary epithelial cells. Journal of Biological Chemistry. 1996;271(21):12414–22.

98. Wu W, Chaudhuri S, Brickley DR, Pang D, Karrison T, Conzen SD. Microarray analysis reveals glucocorticoid-regulated survival genes that are associated with inhibition of apoptosis in breast epithelial cells. Cancer research. 2004;64(5):1757–64.

99. Zeisberg M, Neilson EG. Biomarkers for epithelial-mesenchymal transitions. The Journal of clinical investigation. 2009;119(6):1429–37.

100. Cao X, Cao Y, Li W, Zhang H, Zhu Z. P4HA1/HIF1α feedback loop drives the glycolytic and malignant phenotypes of pancreatic cancer. Biochemical and biophysical research communications. 2019.

101. Xu R. P4HA1 is a new regulator of the HIF-1 pathway in breast cancer. Cell stress. 2019;3(1):27.

102. Gilkes DM, Bajpai S, Chaturvedi P, Wirtz D, Semenza GL. Hypoxia-inducible factor 1 (HIF-1) promotes extracellular matrix remodeling under hypoxic conditions by inducing P4HA1, P4HA2, and PLOD2 expression in fibroblasts. Journal of Biological Chemistry. 2013;288(15):10819–29.

103. Burridge K, Fath K, Kelly T, Nuckolls G, Turner C. Focal adhesions: transmembrane junctions between the extracellular matrix and the cytoskeleton. Annual review of cell biology. 1988;4(1):487–525.

104. Flores-Pérez A, Marchat LA, Rodríguez-Cuevas S, Bautista VP, Fuentes-Mera L, Romero-Zamora D, et al. Suppression of cell migration is promoted by miR-944 through targeting of SIAH1 and PTP4A1 in breast cancer cells. BMC cancer. 2016;16(1):379.

105. Dluzen D, Li G, Tacelosky D, Moreau M, Liu DX. BCL-2 is a downstream target of ATF5 that mediates the prosurvival function of ATF5 in a cell type-dependent manner. Journal of Biological Chemistry. 2011;286(9):7705–13.

106. Greene LA, Lee HY, Angelastro JM. The transcription factor ATF5: role in neurodevelopment and neural tumors. Journal of neurochemistry. 2009;108(1):11–22.

107. Sheng Z, Evans SK, Green MR. An activating transcription factor 5-mediated survival pathway as a target for cancer therapy. Oncotarget. 2010;1(6):457.

108. Miyoshi N, Ishii H, Mimori K, Takatsuno Y, Kim H, Hirose H, et al. Abnormal expression of TRIB3 in colorectal cancer: a novel marker for prognosis. British journal of cancer. 2009;101(10):1664.

109. Dong S, Xia J, Wang H, Sun L, Wu Z, Bin J, et al. Overexpression of TRIB3 promotes angiogenesis in human gastric cancer. Oncology reports. 2016;36(4):2339–48.

110. Helmy N, Prip-Buus C, Vons C, Lenoir V, Abou-Hamdan A, Guedouari-Bounihi H, et al. Oxidation of hydrogen sulfide by human liver mitochondria. Nitric Oxide. 2014;41:105–12.

111. Fagerberg L, Hallström BM, Oksvold P, Kampf C, Djureinovic D, Odeberg J, et al. Analysis of the human tissue-specific expression by genome-wide integration of transcriptomics and antibody-based proteomics. Molecular & Cellular Proteomics. 2014;13(2):397–406.

112. Szabo C, Ransy C, Módis K, Andriamihaja M, Murghes B, Coletta C, et al. Regulation of mitochondrial bioenergetic function by hydrogen sulfide. Part I. Biochemical and physiological mechanisms. British journal of pharmacology. 2014;171(8):2099–122.

113. Gibbins MTG. Metabolic and vascular effects of thiosulfate sulfurtransferase deletion. 2018.

114. Liao A, Xiao Y. Clinic significance of downexpression of LGALS4 gene in colorectal cancer. Cancer Research and Clinic. 2011;23(10):661–3.

115. Rodia MT, Solmi R, Mattia L. TSPAN8 and LGALS4 combination as blood biomarkers for colorectal cancer detection. Cancer Cell & Microenvironment. 2016;3(3).

116. Khanin R, Wit E. How scale-free are biological networks. Journal of computational biology. 2006;13(3):810–8.

